# Large and fast human pyramidal neurons associate with intelligence

**DOI:** 10.1101/296343

**Authors:** Natalia A. Goriounova, Djai B. Heyer, René Wilbers, Matthijs B. Verhoog, Michele Giugliano, Christophe Verbist, Joshua Obermayer, Amber Kerkhofs, Harriët Smeding, Maaike Verberne, Sander Idema, Johannes C. Baayen, Anton W. Pieneman, Christiaan P.J. de Kock, Martin Klein, Huibert D. Mansvelder

## Abstract

It is generally assumed that human intelligence relies on efficient processing by neurons in our brain. Behavioral and brain-imaging studies robustly show that higher intelligence associates with faster reaction times and thicker gray matter in temporal and frontal cortical areas. However, no direct evidence exists that links individual neuron activity and structure to human intelligence. Since a large part of cortical grey matter consists of dendrites, these structures likely determine cortical architecture. In addition, dendrites strongly affect functional properties of neurons, including action potential speed. Thereby, dendritic size and action potential firing may constitute variation in cortical thickness, processing speed, and ultimately IQ.

To investigate this, we took advantage of brain tissue available from neurosurgery and recorded from pyramidal neurons in the medial temporal cortex, an area showing high association between cortical thickness, cortical activity and intelligence. Next, we reconstructed full dendritic structures of recorded neurons and combined these with brain-imaging data and IQ scores from the same subjects. We find that high IQ scores and large temporal cortical thickness associate with larger, more complex dendrites of human pyramidal neurons. We show *in silico* that larger dendrites enable pyramidal neurons to track activity of synaptic inputs with higher temporal precision, due to fast action potential initiation. Finally, we find that human pyramidal neurons of individuals with higher IQ scores sustain faster action potentials during repeated firing. These findings provide first evidence that human intelligence is associated with neuronal complexity, action potential speed and efficient information transfer in cortical neurons.

## Introduction

A fundamental question in neuroscience is what properties of neurons lie at the heart of human intelligence and underlie individual differences in mental ability. Thus far, experimental research on the neurobiological basis of intelligence has largely ignored the neuronal level and has not directly tested what role human neurons play in cognitive ability, mainly due to the inaccessibility of human neurons. Instead, research has either been focused on finding genetic loci that can explain part of the variance in intelligence (Spearman’s *g)* in large cohorts (Lam et al. 2017; Sniekers et al. 2017; Trampush et al. 2017; Coleman et al. 2018) or on identifying brain regions in whole brain imaging studies of which structure or function correlate with IQ scores (Choi et al. 2008; Karama et al. 2009; Hulshoff Pol et al. 2006; Narr et al. 2007; McDaniel 2005; Deary et al. 2010). Some studies have highlighted that variability in brain volume and intelligence may share a common genetic origin (Hulshoff Pol et al. 2006; Posthuma et al. 2002; Sniekers et al. 2017), and individual genes that were identified as associated with IQ scores might aid intelligence by facilitating neuron growth (Sniekers et al. 2017; Coleman et al. 2018) and directly influencing neuronal firing (Lam et al. 2017).

Intelligence is a distributed function that depends on activity of multiple brain regions (Deary et al. 2010). Structural and functional magnetic resonance imaging studies in hundreds of healthy subjects revealed that cortical volume and function of specific areas correlate with *g* (Choi et al. 2008; Karama et al. 2009; Narr et al. 2007). In particular, areas located in the frontal and temporal cortex show a strong correlation of grey matter thickness and functional activation with IQ scores: individuals with high IQ show larger grey matter volume of, for instance, Brodmann areas 21 and 38 (Choi et al. 2008; Deary et al. 2010; Karama et al. 2009; Narr et al. 2007). Cortical grey matter consists for a substantial part of dendrites (Chklovskii et al. 2002; Ikari & Hayashi 1981), which receive and integrate synaptic information and strongly affect functional properties of neurons (Bekkers & Häusser 2007; Eyal et al. 2014; Vetter et al. 2001). Especially high-order association areas in temporal and frontal lobes in humans harbor pyramidal neurons of extraordinary dendritic size and complexity (Elston 2003; Mohan et al. 2015) that may constitute variation in cortical thickness, neuronal function, and ultimately IQ. These neurons and their connections form the principal building blocks for coding, processing, and storage of information in the brain and ultimately give rise to cognition (Salinas & Sejnowski 2001). Given their vast number in the human neocortex, even the slightest change in efficiency of information transfer by neurons may translate into large differences in mental ability. However, whether and how the activity and dendritic structure of single human neurons support human intelligence has not been tested.

To investigate whether structural and functional properties of neurons of the human temporal cortex associate with general intelligence, we collected a unique multimodal data set from human subjects containing single cell physiology, neuronal morphology, pre-surgical MRI scans and IQ test scores (Fig 1). We recorded action potentials (APs) from human pyramidal neurons in superficial layers of temporal cortical tissue resected during neurosurgery (Brodmann areas 38 and 21) and digitally reconstructed their complete dendritic structures. We tested the hypothesis that variation in neuronal morphology can lead to functional differences in AP speed and information transfer and explain variation in IQ scores. In addition to our experimental results, we used computational modelling to understand underlying principles of efficient information transfer in cortical human neurons.

**Figure 1.**
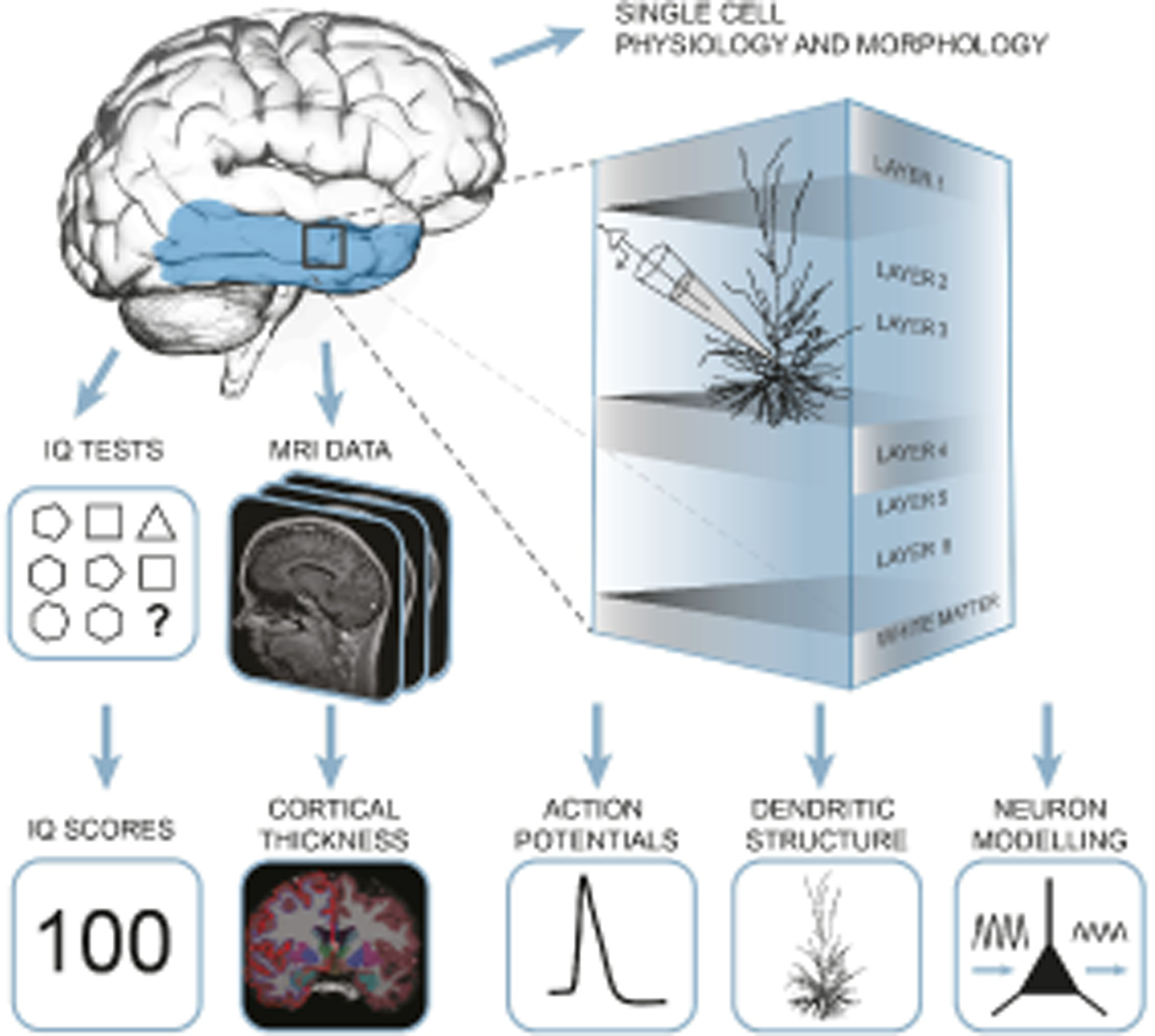
Summary of the approach: multidimensional data set from human subjects contained single cell physiology, neuronal morphology, MRI and IQ test scores (WAIS FSIQ). The area of the brain highlighted in blue indicates the location of cortical thickness measurements, black square indicates the typical origin of resected cortical tissue.

## Results

### 1. IQ scores positively correlate with cortical thickness of the temporal lobe

Cortical thickness of the temporal lobe has been associated with IQ scores in hundreds of healthy subjects (Choi et al. 2008; Deary et al. 2010; Hulshoff Pol et al. 2006; Karama et al. 2009; Narr et al. 2007) and we first asked whether this applies to the subjects in our study as well. From T1-weighted MRI scans obtained prior to surgery, we determined temporal cortical thickness in 35 subjects using voxel-based morphometry. We selected temporal cortical areas corresponding to surgical resections (Fig 2a) and collapsed the measurements for temporal lobes to one mean value for cortical thickness for each subject. In line with previous studies (Choi et al. 2008; Deary et al. 2010; Hulshoff Pol et al. 2006; Narr et al. 2007; Karama et al. 2009), mean cortical thickness in temporal lobes positively correlated with IQ scores of the subjects (Fig 2b).

**Figure 2.**
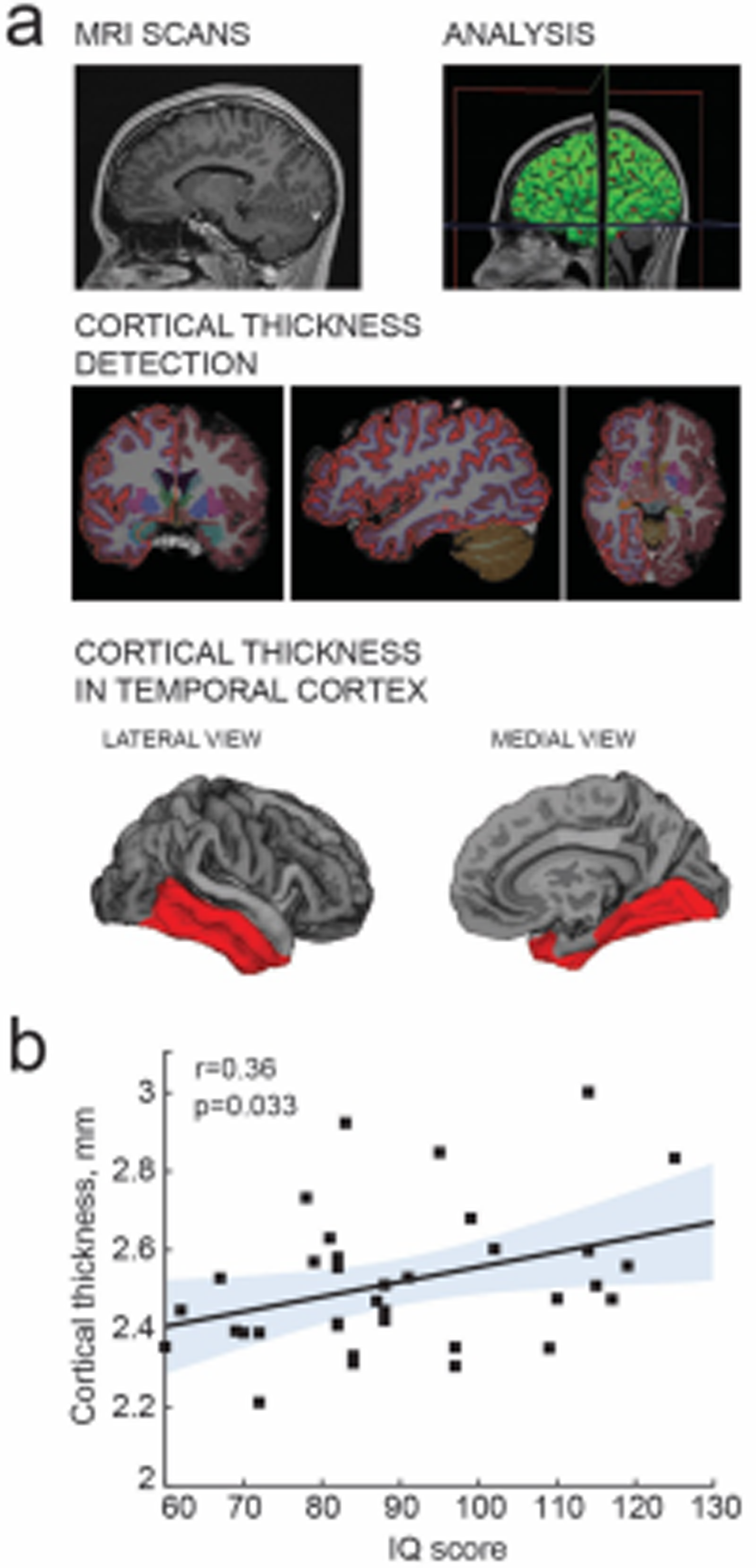
IQ scores positively correlate with cortical thickness of the temporal lobe. (**a**) MRI analysis pipeline: 1) Presurgical MRI T1-weighted scans; 2) Morphometric analysis; 3) Detection of cortical thickness from pial and white-grey matter boundaries; 4) selection of temporal cortical area for correlations with IQ in B (highlighted in red). (**b**) Average cortical thickness in temporal lobe (from area highlighted in red in A4) positively correlates with IQ scores form the same subjects (n subjects=35). Here and in figures below, Pearson correlation coefficients and p-values are reported in graph insets, the solid line represents linear regression (R^2^=0.13), shaded area indicates 95% confidence bounds of the fit.

### 2. IQ scores positively correlate with dendritic structure of temporal cortical pyramidal cells

Cortical association areas in temporal lobes play a key role in high-level integrative neuronal processes and its superficial layers harbor neurons of increased neuronal complexity (DeFelipe et al. 2002; Elston 2003; Scholtens et al. 2014; van den Heuvel et al. 2015). In rodents, the neuropil of cortical association areas consists for over 30% of dendritic structures (Ikari & Hayashi 1981). To test the hypothesis that human temporal cortical thickness is associated with dendrite size, we used 72 full reconstructions of biocytin-labelled temporal cortical pyramidal neurons from human layer 2, 3 and 4 part of which was previously reported (Mohan et al. 2015). Surgically obtained cortical tissue was non-pathological and was resected to gain access to deeper structures containing disease focus (Mohan et al. 2015; Testa-Silva et al. 2014; Testa-Silva et al. 2010; Verhoog et al. 2016; Verhoog et al. 2013) (typically medial temporal sclerosis or hippocampal tumor; Suppl. Table 1). In line with the non-pathological status of tissue, we observed no correlations of cellular parameters or IQ scores with the subject’s disease history and age (Suppl Figs 1&2). We calculated total dendritic length (TDL) for each neuron and mean TDL from multiple cells for each subject and correlated these mean TDL values to mean temporal cortical thickness from the same subject. We found that dendritic length positively correlated with mean temporal lobe cortical thickness, indicating that dendritic structure of individual neurons contributes to the overall cytoarchitecture of temporal cortex (Fig 3a).

**Figure 3.**
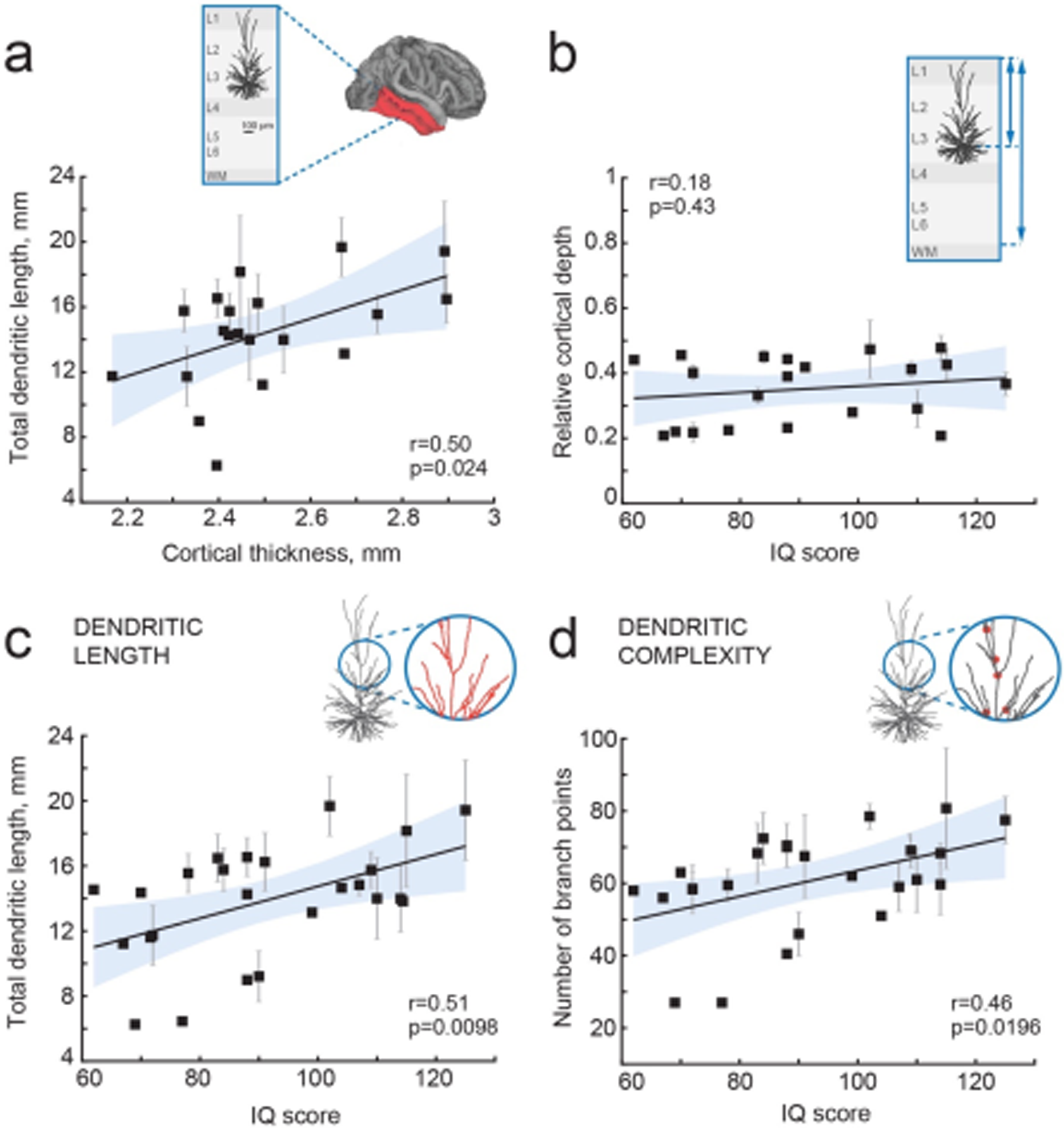
IQ scores positively correlate with dendritic structure of temporal cortical pyramidal cells. (**a**) Average total dendritic length in pyramidal cells in superficial layers of temporal cortex positively correlates with cortical thickness in temporal lobe from the same hemisphere (area shaded in A, n subjects=20; n cell=57, R^2^=0.25). Inset shows a scheme of cortical tissue with a digitally reconstructed neuron and the brain area for cortical thickness estimation (red) (**b**) Cortical depth of pyramidal neurons, relative to cortical thickness in temporal cortex from the same hemisphere, does not correlate with IQ score (n subjects=21, R^2^=0.03). Inset represent the cortical tissue, blue lines indicate the depth of neuron and cortical thickness (**c**) Total dendritic length (TDL) and (**d**) number of dendritic branch points positively correlate with IQ scores from the same individuals (n subjects=25, n cells=72, TDL R^2^=0.26, Branch points R^2^=0.22). Data are mean per subject ± SEM.

TDL is in part determined by the soma location within cortical layers: cell bodies of pyramidal neurons with larger dendrites typically lie deeper, at larger distance from pia (Mohan et al. 2015). To exclude a systematic bias in sampling, we determined the cortical depth of each neuron relative to the subject’s temporal cortical thickness in the same hemisphere. There was no correlation between IQ score and relative cortical depth of pyramidal neurons indicating that we sampled neurons at similar depths across subjects (Fig 3b). Finally, we tested whether mean TDL and complexity of pyramidal neurons relates to subjects’ IQ scores. We found a strong positive correlation between individual’s pyramidal neuron TDL and IQ scores (Fig 3c) as well as between number of dendritic branch points and IQ scores (Fig 3d). Thus, larger and more complex pyramidal neurons in temporal association area may partly contribute to thicker cortex and link to higher intelligence.

### 3. Larger dendrites lead to faster AP onset and improved encoding properties

Dendrites not only receive most synapses in neurons, but dendritic morphology and ionic conductances act in concert to regulate neuronal excitability (Bekkers & Häusser 2007; Eyal et al. 2014; Vetter et al. 2001). Increase in the size of the dendritic compartment was shown *in silico* to accelerate APs and improve encoding capability of simplified model neurons (Eyal et al. 2014). Further, human neocortical pyramidal neurons, which are three times larger than rodent pyramidal neurons (Mohan et al. 2015), have faster AP onsets compared to rodent neurons and are able to track and encode fast synaptic inputs and sub-threshold changes in membrane potential with high temporal precision (Testa-Silva et al. 2014). We asked whether the observed differences in TDL between human pyramidal neurons affected their encoding properties and ability to transfer information. To this end, we incorporated the 3-dimensional dendritic reconstructions of the human pyramidal neurons into *in silico* models and equipped them with excitable properties (see Supplementary Methods). We first simulated somatic APs in model neurons. We found that TDL of model neurons positively correlated with the steepness of AP onsets (Fig. 4 a,b) and larger dendrites enable neurons to generate faster APs.

**Figure 4.**
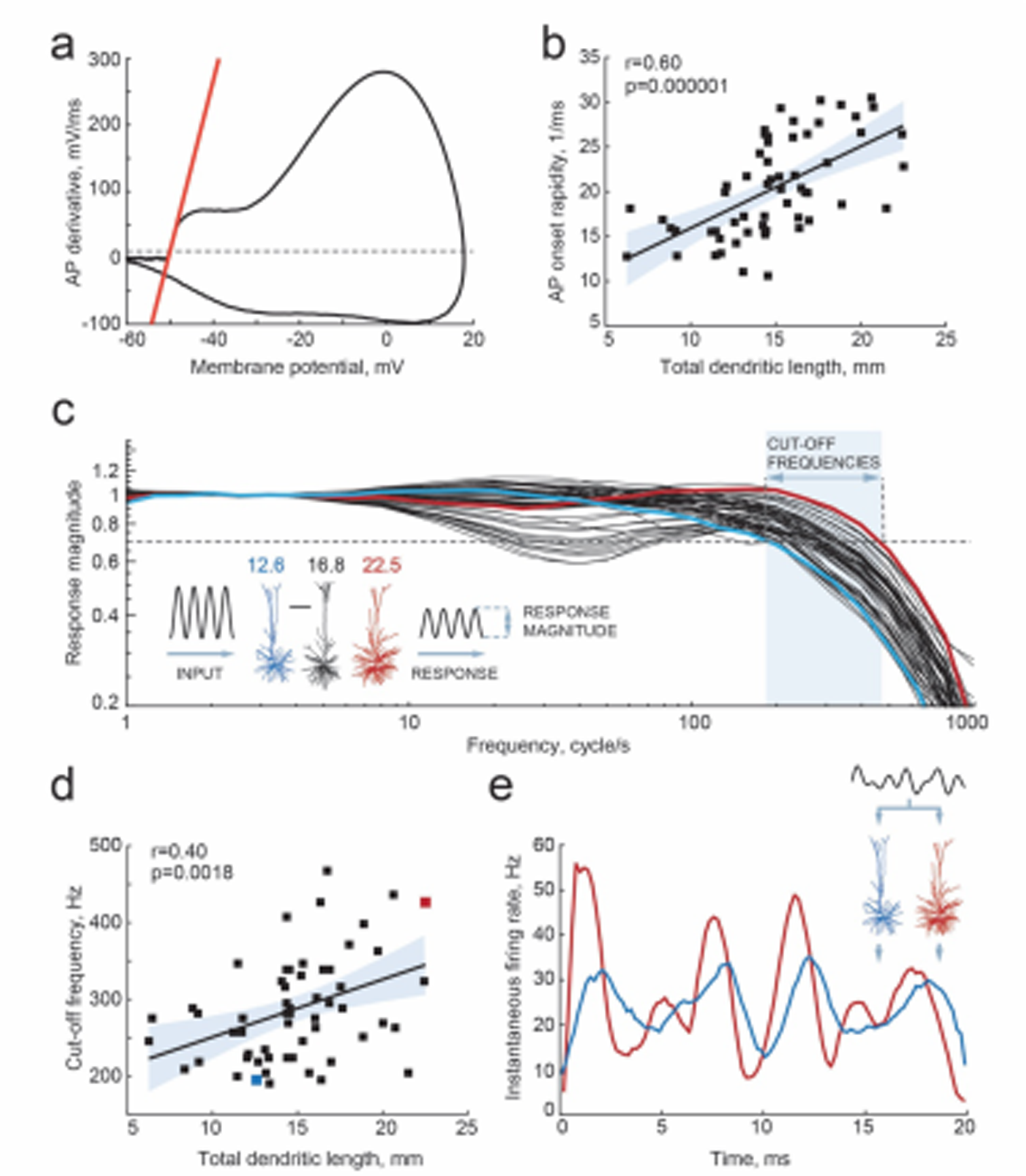
Larger dendrites lead to faster AP onset and improved encoding properties. (**a,b**) Higher TDL results in faster onsets of model-generated APs: **(a)** example phase plot of an AP is shown with a red line representing onset rapidity - slope of AP derivative at 10 mV/ms (grey dashed line)**; (b)** onset rapidity values of simulated APs positively correlate with TDL (R^2^=0.36). **(c)** Model neurons received simulated sinusoidal current-clamp inputs and generated spiking responses of different magnitudes and frequencies. Red and blue traces are response magnitudes of example neurons with low (blue) and large (red) TDLs; inset shows examples of morphological reconstructions with their TDLs in mm shown above. Cut-off frequencies are defined within the frequency range (shaded area) at which model neuron can still track the inputs reliably (produce response of 0.7 response magnitude, dashed line). (**d**) Cut-off frequencies positively correlate with TDL (R^2^=0.16; example neurons from panel (**c**) are highlighted by the same colors). (**e**) Responses to the same input in two example neurons from panels **(b)** and **(c):** instantaneous firing frequency of the model neuron with large TDL (red) follows the input with higher temporal precision than the model neuron with smaller TDL (blue).

The exact timing of action potential firing allows cortical neurons to pass on temporal information provided by synaptic inputs (Köndgen et al. 2008; Ilin et al. 2013; Testa-Silva et al. 2014). Single pyramidal neurons do not sustain high frequency firing and generally do not encode high frequency synaptic input content in rate coding. Instead, the precision in timing of AP initiation does allow these neurons to encode incoming high frequency information in their output. In contrast to rodent neurons, human neurons can encode sub-threshold membrane potential changes on a sub-millisecond timescale by timing of APs (Testa-Silva et al. 2014). This synaptic input tracking capacity strongly relies on the speed of AP generation (Ilin et al. 2013). Faster APs allow neurons to respond to fast synaptic inputs, which will be missed if AP generation is too slow. Thereby, neurons with faster APs can translate higher frequencies of synaptic membrane potential fluctuations into AP timing and ultimately encode more information. Although ‘ball-and-stick’ models suggest that neurons with larger dendritic compartments have faster APs and can encode more information in their output (Eyal et al., 2014), it is not known whether this holds true for the human cortical pyramidal neurons we recorded from. We tested this by simulating sinusoidal current inputs of increasing frequencies into in silico representations of the neurons we recorded and reconstructed, and studied how the timing of AP firing of these neurons followed sub-threshold membrane potential changes. We find that human neurons with larger TDL can reliably time their APs to faster membrane potential changes, with cut-off frequencies up to 400-500 Hz, while smaller neurons had their cut-off frequencies already at 200 Hz (Fig 4c, d). Furthermore, there was a significant positive correlation between the dendritic length and the cut-off frequency (Fig. 4d). Finally, given the same input - composed of the sum of three sinusoids of increasing frequencies - larger neurons were able to better encode rapidly changing temporal information into timing of AP firing, compared to smaller neurons (Fig. 4e). Thus, we find that differences in dendritic length of human neurons lead to faster APs and thereby to wider frequency bandwidths of encoding synaptic inputs into timing of AP output.

### 4. Higher IQ scores associate with faster APs

Since cortical pyramidal neurons with large dendrites have faster APs and can encode more information in AP output, and since large dendrites also associate with higher IQ scores, we next asked whether human cortical pyramidal neurons from individuals with higher IQ scores generate faster APs. To test this, we made whole-cell recordings from pyramidal cells in acute slices of temporal cortex (31 subjects, 129 cells, Fig 5) and recorded APs at different firing frequencies in response to depolarizing current steps. The maximum rise speed of APs depended on the firing history of the cell, with the first AP in the train having the highest AP rise speed and slowing down with increasing instantaneous firing frequency, the time interval between subsequent APs (Fig 5b-d). To test whether AP rise speeds differed between IQ groups, we split all AP rise speed data into two groups based on IQ score – above and below 100. Although the AP rise speed of the first AP was not different between high and low IQ groups (Fig 5c), the AP slowed down stronger in individuals with IQ scores below 100 compared to APs of individuals with IQ scores above 100 (Fig 5d). At higher instantaneous firing frequencies (20-40 Hz), the AP rise speed was higher in individuals with IQ scores above 100 (Fig 5c right; AP rise speed high IQ=338.4±26.03 mV/ms; AP rise speed low IQ=268.1±12.20 mV/ms, t-test p=0.0113). We next calculated the slowing of APs with increasing instantaneous frequency by normalizing rise speeds of APs to the rise speed of the first AP in the train. Relative to first AP, rise speeds at 20-40 Hz showed significant slowing in subjects with IQ scores below 100 and decreased to 74% of the initial AP rise speed. In contrast, in neurons from individuals with IQ scores above 100, AP rise speed remained on average at 84% (Fig 5d right, high IQ=0.84±0.014; low IQ=0.74±0.024, t-test p value=0.037).

**Figure 5.**
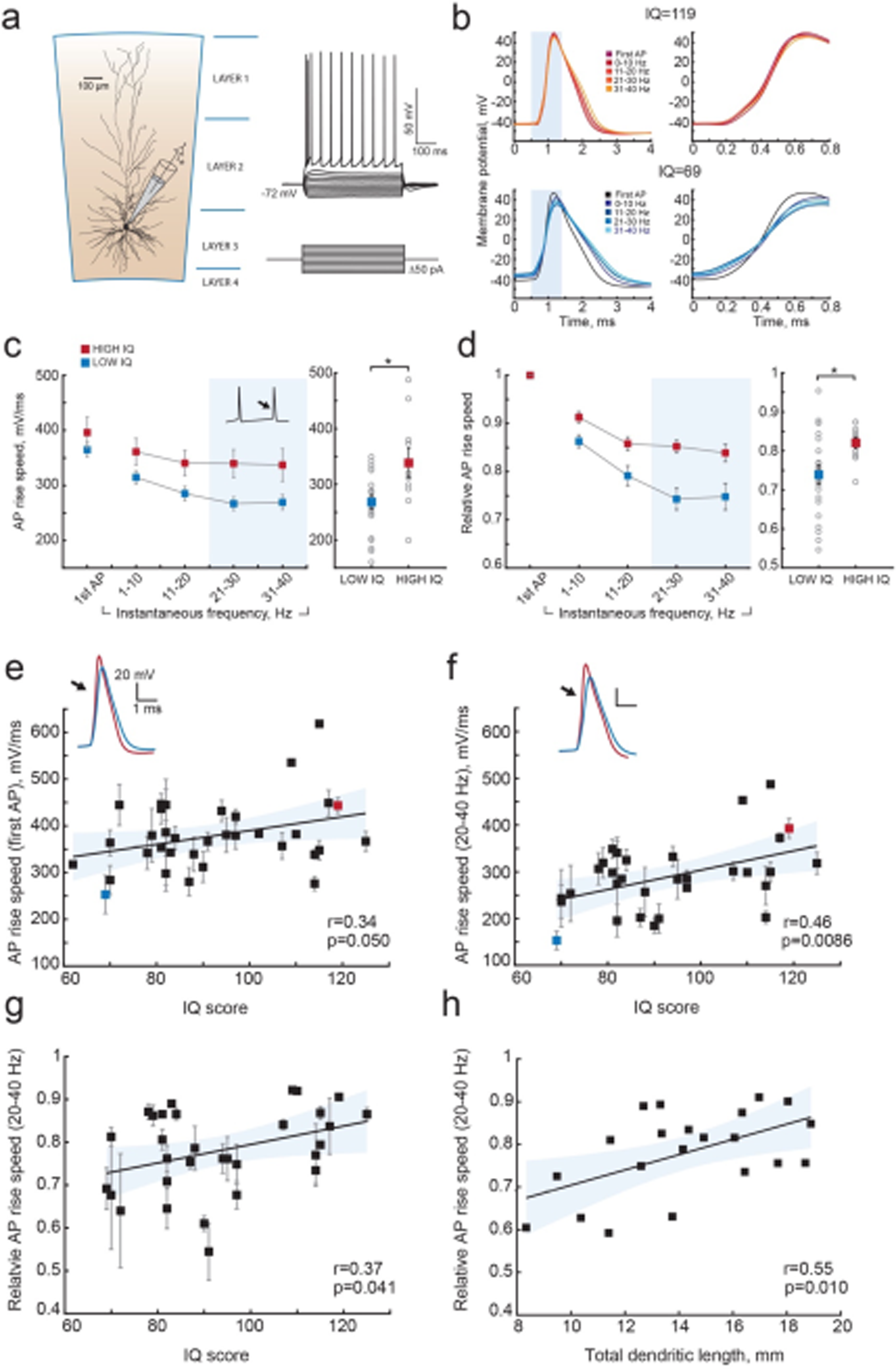
Higher IQ scores associate with faster AP initiation. (**a**) Scheme of a whole-cell recording showing biocytin reconstruction of a pyramidal neuron from human temporal cortex. Right: typical voltage responses to depolarizing somatic current injections. (**b**) Examples of AP traces at increasing instantaneous firing frequencies (frequency is shown in color code in insets) recorded from a subject with IQ=119 (above panel, red) and a subject with IQ=69 (lower panel, blue). AP rising phase in shaded area is displayed to the right (**c**) APs from subjects with higher IQ are better able to maintain their rise speed at increasing frequencies. Average (per neuron and subject) AP rise speeds and (**d**) relative to first AP rise speeds in neurons from subjects with IQ<100 (red, n subjects=10, n cells=38) and subjects with IQ>100 (blue, n subjects=21, n cells=91) are displayed against instantaneous firing frequency. Right: data points in shaded area are shown as averaged values for 20-40 Hz (filled squares are group means, open circles are mean rise speeds per subject), *p<0.05. (**e**) IQ scores positively correlate to the rise speeds of first AP in the train (n subjects 31, n cells=129; R^2^=0.16), (**f**) AP rise speeds at 20-40 Hz (same data as right panel in (c), R^2^=0.21) and (**g**) relative AP rise speeds at 20-40 Hz (same as right panel in (d), R^2^=0.14). (**h**) Larger neurons show less slowing of AP rise speed at higher frequencies: relative AP rise speeds at 20-40 Hz for individual neurons are plotted as a function of their TDL (n=21 cells, R^2^=0.30).

We further investigated whether these differences at the group level reflected correlations between individual IQ scores and AP rise speeds. We correlated mean AP rise speeds both of the first AP and AP at 20-40 Hz from all neurons of the same subject to the subjects IQ score. While the AP rise speed of the first AP in the train showed only a moderate correlation (Fig 5e), AP rise speeds at instantaneous frequencies of 20-40 Hz strongly correlated with IQ scores (Fig 5f). Importantly, also relative to first AP values show significant positive correlations with IQ, indicating that it is the relative slowing of APs that associates with intelligence (Fig 5g). Finally, we asked whether the slowing of APs relates to the dendritic size of the same neurons, as our model results suggest. We find that larger neurons show less slowing of AP rise speed (higher relative AP speeds) at 20-40 Hz. These findings reveal that higher IQ scores are accompanied by faster APs during repeated AP firing, while lower IQ scores associate with increased AP fatigue during elevated neuronal activity. Thus, neurons from individuals with higher IQ scores are better equipped to process synaptic signals at high rates and at faster time scales, which is necessary to encode large amounts of information accurately and efficiently.

## Discussion

Our findings provide a first insight into the possible cellular nature of human intelligence and explain individual variation in IQ scores based on neuronal properties: faster AP rise speed during neuronal activity and more complex, extended dendrites associate with higher intelligence. AP kinetics have profound consequences for information processing. In vivo, neurons are constantly bombarded by high frequency synaptic inputs and the capacity of neurons to keep track and phase-lock to these inputs determines how much of this synaptic information can be passed on to other neurons (Testa-Silva et al. 2014). The brain operates at a millisecond time-scale and even sub-millisecond details of spike trains contain behaviorally relevant information that can steer behavioral responses (Nemenman et al. 2008). Indeed, one of the most robust and replicable findings in behavioral psychology is the association of intelligence scores with measures of cognitive information-processing speed (Barrett et al. 1986). Specifically, reaction times (RT) in simple RT tasks provide a better prediction of IQ than other speed-of-processing tests, with a regression coefficient of 0.447 (Vernon 1983). In addition, high positive correlations between RT and other speed-of-processing tests suggest the existence of a common mental speed factor (Vernon 1983). Recently, these classic findings were confirmed in a large longitudinal population-based study counting more than 2000 participants (Der & Deary 2017). Especially strong correlations between RT and general intelligence were reported for a slightly more complex 4-choice (Der & Deary 2017). Our results provide a biological cellular explanation for such mental speed factors: in conditions of increased mental activity or more demanding cognitive task, neurons of individuals with higher IQ are able to sustain fast action potentials and can transfer more information content from synaptic input to AP output.

Pyramidal cells are integrators and accumulators of synaptic information. Larger dendrites can physically contain more synaptic contacts and integrate more information. Indeed, human pyramidal neuron dendrites receive twice as many synapses as in rodents (DeFelipe et al. 2002) and cortico-cortical whole-brain connectivity positively correlates with the size of dendrites in these cells (Scholtens et al. 2014; van den Heuvel et al. 2015). A gradient in complexity of pyramidal cells in cortical superficial layers accompanies the increasing integration capacity of cortical areas, indicating that larger dendrites are required for higher-order cortical processing (Elston 2003; van den Heuvel et al. 2015). Our results align well with these findings, suggesting that the neuronal complexity gradient also exists from individual to individual and could explain differences in mental ability.

Within human cortex, association areas contain neurons with larger and more complex dendrites than primary sensory areas, while neuronal cell body density is lower in cortical association areas compared to primary sensory areas (Elston 2003; DeFelipe et al. 2002). Larger neurons are not as tightly packed together within cortical space as smaller cells. A recent study by Genc et al (2018) used multi-shell diffusion tensor imaging to estimate parieto-frontal cortical dendritic density and found that higher IQ scores correlated with lower values of dendritic density (Genç et al. 2018). This may indicate that parietofrontal cortical areas in individuals with higher IQ scores have less densely packed neurons, and may suggest that these neurons are larger. In our study, we carefully determined the amount and complexity of dendrite for each neuron, being a computational unit within the cortex with well-defined input-output signals. Taking the results of Genc et al (2018) and our study together may suggest that the neuronal circuitry associated with higher intelligence is organized in a sparse and efficient manner, where larger and more complex pyramidal cells occupy larger cortical volume.

Larger dendrites have an impact on excitability of cells (Bekkers & Häusser 2007; Vetter et al. 2001) and determine the shape and rapidity of APs (Eyal et al. 2014). Increasing the size of dendritic compartments *in silico* lead to acceleration of AP onset and increased encoding capability of neurons (Eyal et al. 2014). Both in models and slice recordings, changes of AP initiation dynamics were shown to fundamentally modify encoding of fast changing signals and the speed of communication between ensembles of cortical neurons (Eyal et al. 2014; Ilin et al. 2013). Neurons with fast AP onsets can encode high frequencies and respond quickly to subtle input changes. However, this ability can be impaired and response speed is decreased when AP onsets are slowed down by experimental manipulations (Ilin et al. 2013). Our results not only demonstrate that AP speed depends on dendritic length and influences information transfer, but also show that both dendritic length and AP speed in human neurons correlate with intelligence. Thus, individuals with larger dendrites are better equipped to transfer synaptic information at higher frequencies.

Remarkably, dendritic morphology and different parameters of AP waveform are also parameters that we have previously identified as showing pronounced differences between humans and other species (Mohan et al. 2015; Testa-Silva et al. 2014). Human pyramidal cells in layers 2/3 have 3-fold larger and more complex dendrites than in macaque or mouse (Mohan et al. 2015). Moreover, human APs have lower firing threshold and faster AP onset kinetics both in single APs and during repeated firing (Testa-Silva et al. 2014). These differences across species may suggest evolutionary pressure on both dendritic structure and AP waveform and emphasize specific adaptations of human pyramidal cells in association areas for cognitive functions.

Our results were obtained from patients undergoing neurosurgical procedure and, thus, may potentially raise questions on how representative our findings are for normal healthy human subjects. Although no healthy controls can be used for single cell measurements, we addressed this issue in the following way. Firstly, in all patients, the resected neocortical tissue was not part of epileptic focus or tumor and displayed no structural or functional abnormalities in preoperative MRI, electrophysiological recordings or microscopic investigation of stained tissue. Secondly, none of the parameters correlated with age at epilepsy onset, seizure frequency, age or disease duration (Suppl Fig. 1). Thirdly, IQ, dendritic length or AP rise speed were not different across different patient groups (Suppl Fig. 2). Finally, the cortical thickness correlation with general intelligence we observe in our study was also reported in hundreds of healthy subjects. Taken together, these results indicate that our findings are not likely to be influenced by disease background of the subjects.

In conclusion, our results provide first evidence that already at the level of individual neurons, such parameters as dendritic size and ability to maintain fast responses link to general mental ability. Multiplied by an astronomical number of cortical neurons in our brain, very small changes in these parameters may lead to large differences in encoding capabilities and information transfer in cortical networks and result in a speed advantage in mental processing and, finally, in faster reaction times and higher cognitive ability.

## Methods

### Human subjects and brain tissue

All procedures were performed with the approval of the Medical Ethical Committee of the VU University Medical Centre, and in accordance with Dutch licence procedures and the Declaration of Helsinki. Written informed consent was provided by all subjects for data and tissue use for scientific research. All data were anonymized.

Human cortical brain tissue was removed as a part of surgical treatment of the subject in order to get access to a disease focus in deeper brain structures (hippocampus or amygdala) and typically originated from gyrus temporalis medium (Brodmann areas 21 or 38, occasionally gyrus temporalis inferior or gyrus temporalis superior). Speech areas were avoided during resection surgery through functional mapping. We obtained neocortical tissue from 44 patients (24 females, 20 males; age range 18–66 years, Supplementary table 1) treated for mesial temporal sclerosis, removal of a hippocampal tumor, low grade hippocampal lesion, cavernoma or another unspecified temporal lobe pathology. From 35 of these patients we also obtained pre-surgical MRI scans, from 31 patients we recorded Action Potentials from 129 cells and from 25 patients we had fully reconstructed dendritic morphologies from 72 cells.

In all patients, the resected neocortical tissue was not part of epileptic focus or tumor and displayed no structural/functional abnormalities in preoperative MRI investigation, electrophysiological whole-cell recordings or microscopic investigation of stained tissue. The physiological recordings, subsequent morphological reconstructions, morphological and action potential analysis were performed blind to the IQ of the patients.

### IQ scores

Total IQ scores were obtained from all 44 subjects using the Dutch version of Wechsler Adult Intelligence Scale-III (WAIS-III) and in some cases WAIS-IV and consisted of following subtests: information, similarities, vocabulary, comprehension, block design, matrix reasoning, visual puzzles, picture comprehension, figure weights, digit span, arithmetic, symbol search and coding.

The tests were performed as a part of neuropsychological examination shortly before surgery, typically within one week.

### MRI data and cortical thickness estimation

T1-weighted brain images (1 mm thickness) were acquired with a 3T MR system (Signa HDXt, General Electric, Milwaukee, Wisconsin) as a part of pre-surgical assessment (number of slices=170-180). Cortical reconstruction and volumetric segmentation was performed with the Freesurfer image analysis suite (http://freesurfer.net) (Fischl & Dale 2000). The processing included motion correction, transformation to the Talairach frame. Calculation of the cortical thickness was done as the closest distance from the grey/white boundary to the grey/CSF boundary at each vertex and was based both on intensity and continuity information from the entire three-dimensional MR volume (Fischl & Dale 2000). Neuroanatomical labels were automatically assigned to brain areas based on Destrieux cortical atlas parcellation as described in (Fischl 2004). For averaging, the regions in temporal lobes were selected based on Destrieux cortical atlas parcellation in each subject.

### Slice preparation

Upon surgical resection, the cortical tissue block was immediately transferred to ice-cold artificial cerebral spinal fluid (aCSF) containing in (mM): 110 choline chloride, 26 NaHCO3, 10 D-glucose, 11.6 sodium ascorbate, 7 MgCl2, 3.1 sodium pyruvate, 2.5 KCl, 1.25 NaH2PO4, and 0.5 CaCl2 (300 mOsm) and transported to the neurophysiology laboratory (within 500 m from the operating room). The transition time between resection of the tissue and the start of preparing slices was less than 15 minutes. After removing the pia and identifying the pia-white matter axis, neocortical slices (350 mm thickness) were prepared in ice-cold slicing solution (same composition as described above). Slices were then transferred to holding chambers in which they were stored for 30 minutes at 34 C^o^ and for 30 minutes at room temperature before recording in aCSF, which contained (in mM): 126 NaCl; 3 KCl; 1 NaH2PO4; 1 MgSO4; 2 CaCl2; 26 NaHCO3; 10 glucose (300 mOsm), bubbled with carbogen gas (95% O2/5% CO2), as described previously (Mohan et al. 2015; Testa-Silva et al. 2014; Testa-Silva et al. 2010; Verhoog et al. 2013; Verhoog et al. 2016).

### Electrophysiological recordings

Cortical slices were visualized using infrared differential interference contrast (IR-DIC) microscopy. After the whole cell configuration was established, membrane potential responses to steps of current injection (step size 30-50 pA) were recorded. None of the neurons showed spontaneous epileptiform spiking activity. Recordings were made using Multiclamp 700A/B amplifiers (Axon Instruments) sampling at frequencies of 10 to 50 kHz, and lowpass filtered at 10 to 30 kHz. Recordings were digitized by pClamp software (Axon) and later analyzed off-line using custom-written Matlab scripts (MathWorks). Patch pipettes (3–5 MOhms) were pulled from standard-wall borosilicate capillaries and filled with intracellular solution containing (in mM): 110 K-gluconate; 10 KCl; 10 HEPES; 10 K-phosphocreatine; 4 ATP-Mg; 0.4 GTP, pH adjusted to 7.2–7.3 with KOH; 285–290 mOsm, 0.5 mg/ml biocytin. All experiments were performed at 32C^o^–35C^o^. Only cells with bridge balance of <15 MOhm were used for further analysis.

### Morphological analysis

During electrophysiological recordings, cells were loaded with biocytin through the recording pipette. After the recordings the slices were fixed in 4% paraformaldehyde and the recorded cells were revealed with the chromogen 3,3-diaminobenzidine (DAB) tetrahydrochloride using the avidin-biotin–peroxidase method (Horikawa & Armstrong 1988). Slices were mounted on slides and embedded in mowiol (Clariant GmbH, Frankfurt am Main, Germany). Neurons without apparent slicing artifacts and uniform biocytin signal were digitally reconstructed using Neurolucida software (Microbrightfield, Williston, VT, USA), using a *100 oil objective. After reconstruction, morphologies were checked for accurate reconstruction in x/y/z planes, dendritic diameter, and continuity of dendrites. Finally, reconstructions were checked using an overlay in Adobe Illustrator between the Neurolucida reconstruction and Z-stack projection image from Surveyor Software (Chromaphor, Oberhausen, Germany).

Superficial layers pyramidal neurons were identified based on morphological and electrophysiological criteria at cortical depth within 400-1400 μm from cortical surface, that we previously found to correspond to cortical layers 2, 3 and 4 in humans (Mohan et al. 2015). For each neuron, we extracted total dendritic length (TDL) and number of branch points and computed average TDL and average number of branch points for each subject by pulling data from all cells from one subject (1 to 8 cells per subject).

### Neuronal modelling

We constructed multicompartmental spiking neuronal models of human pyramidal neurons for each the digitally reconstructed 3-dimensional morphologies considered in this study. Models were simulated using NEURON (Carnevale & Hines 2006). As in the experiments of Köndgen *et al.* (Köndgen et al. 2008) and Testa-Silva *et al.* (Testa-Silva et al. 2014), we probed the dynamical transfer properties of each model neuron. We injected a sinusoidally oscillating input current in the soma, allowing us to temporally modulate the instantaneous output firing rate of each model neuron and quantify its output ‘transfer gain’ (Fig. 4). When studied in this way, the transfer properties of model neurons resemble those of electronic filters, whose low-pass performances in the Fourier domain define how fast they can follow input changes (for more detailed information see Supplementary Methods).

### Action Potential waveform analysis

Action Potential (AP) waveforms were extracted from voltage traces recorded in response to intracellular current injections and sorted according to their instantaneous firing frequency. Instantaneous frequency was determined as 1/time to previous AP. Subsequently all APs were binned in 10Hz bins, while first APs in each trace were isolated in a separate bin.

AP rise speed was defined as the peak of AP derivative (dV/dt). For each analyzed cell, representative APs with all parameters were plotted for visual check to avoid errors in the analysis.

For each neuron, the mean values of AP rise speed in a given frequency bin were obtained by averaging all APs within that frequency bin. Relative AP rise speeds were calculated by dividing the mean AP rise speed in each frequency bin (1-10 Hz, 11-20 Hz, 21-30 Hz and 31 to 40 Hz) by the mean first AP rise speed (first APs in the train of APs).

To obtain AP values for each subject, AP parameters within each frequency bin were averaged for all neurons from one subject.

### Statistical analysis

Statistical significance of all correlations between parameters was determined using Pearson correlation and linear regression (using Matlab, version R2017a, Mathworks). As multiple cells were measured per subject, correlations were calculated on mean parameter values per subject. All Pearson correlation coefficients and p values for correlations are shown in figure insets, R^2^ coefficients and sample sizes are shown in figure legends.

For statistical analysis of AP data, we divided all subjects according to their IQ into 2 groups: group with IQ>100 and a group with IQ<100. Differences between 2 IQ groups in AP rise times were statistically tested using Student t-test. For analysis of different patient groups (Fig S2) ANOVA test was applied for each parameter separately.

## Acknowledgments

We thank dr. Linda Douw for her assistance with the analysis of brain imaging data and Mr. M. Wijnants for his technical assistance and to the supercomputer facilities CalUA (University of Antwerp) for computing time. N.A.G. received funding for this work from the from the Netherlands Organization for Scientific Research (NWO; VENI grant). H.D.M. received funding for this work from the Netherlands Organization for Scientific Research (NWO; VICI grant), ERC StG “BrainSignals”, and EU H2020 “Human Brain Project” grant agreement no. 604102.

M. G. has received funding from EU H2020 “Human Brain Project” no. 720270, and the Flemish Research Foundation (grant no. G0F1517N).

## Author contributions

NAG: performed AP recordings; analysed data; conceived study design; wrote manuscript

D.H contributed to data analysis

R.W. contributed to data analysis

MBV, performed AP recordings and contributed to data analysis

M.G. performed computational modelling

C.V. performed computational modelling JO and AK: performed AP recordings HS and MV: contributed IQ data

SI, JCB: performed neurosurgery and provided MRI data

AP; contributed morphology data CPJdeK: contributed morphology data

MK: contributed IQ data and study design

HDM: conceived study design, wrote manuscript

## References

Barrett, P., Eysenck, H.J. & Lucking, S., 1986. Reaction time and intelligence: A replicated study. Intelligence, 10(1), pp.9–40.

Bekkers, J.M. & Häusser, M., 2007. Targeted dendrotomy reveals active and passive contributions of the dendritic tree to synaptic integration and neuronal output. Proceedings of the National Academy of Sciences of the United States of America, 104(27), pp.11447–11452.

Carnevale, N.T. & Hines, M.L., 2006. The NEURON Book, Cambridge: Cambridge University Press.

Chklovskii, D.B., Schikorski, T. & Stevens, C.F., 2002. Wiring optimization in cortical circuits. Neuron, 34(3), pp.341–347.

Choi, Y.Y. et al., 2008. Multiple bases of human intelligence revealed by cortical thickness and neural activation. The Journal of neuroscience, 28(41), pp.10323–10329.

Coleman, J.R.I. et al., 2018. Biological annotation of genetic loci associated with intelligence in a meta-analysis of 87,740 individuals. Molecular psychiatry, 533, p.533.

Deary, I.J., Penke, L. & Johnson, W., 2010. The neuroscience of human intelligence differences. Nature reviews. Neuroscience, 11(3), pp.201–211.

DeFelipe, J., Alonso-Nanclares, L. & Arellano, J.I., 2002. Microstructure of the neocortex: comparative aspects. Journal of neurocytology, 31(3-5), pp.299–316.

Der, G. & Deary, I.J., 2017. The relationship between intelligence and reaction time varies with age: Results from three representative narrow-age age cohorts at 30, 50 and 69 years. Intelligence, 64, pp.89–97.

Elston, G.N., 2003. Cortex, cognition and the cell: new insights into the pyramidal neuron and prefrontal function. Cerebral Cortex.

Eyal, G. et al., 2014. Dendrites impact the encoding capabilities of the axon. The Journal of neuroscience, 34(24), pp.8063–8071.

Fischl, B., 2004. Automatically Parcellating the Human Cerebral Cortex. Cerebral cortex, 14(1), pp.11–22.

Fischl, B. & Dale, A.M., 2000. Measuring the thickness of the human cerebral cortex from magnetic resonance images. Proceedings of the National Academy of Sciences of the United States of America, 97(20), pp. 11050–11055.

Geng, E. et al., 2018. Diffusion markers of dendritic density and arborization in gray matter predict differences in intelligence. Nature communications, 9(1), p.1905.

Horikawa, K. & Armstrong, W.E., 1988. A versatile means of intracellular labeling: injection of biocytin and its detection with avidin conjugates. Journal of Neuroscience Methods, 25(1), pp.1–11.

Hulshoff Pol, H.E. et al., 2006. Genetic contributions to human brain morphology and intelligence. The Journal of neuroscience, 26(40), pp.10235–10242.

Ikari, K. & Hayashi, M., 1981. Aging in the neuropil of cerebral cortex--a quantitative ultrastructural study. Folia psychiatrica et neurologica japonica, 35(4), pp.477–486.

Ilin, V. et al., 2013. Fast computations in cortical ensembles require rapid initiation of action potentials. The Journal of neuroscience, 33(6), pp.2281–2292.

Karama, S. et al., 2009. Positive association between cognitive ability and cortical thickness in a representative US sample of healthy 6 to 18 year-olds. Intelligence, 37(2), pp.145–155.

Köndgen, H. et al., 2008. The dynamical response properties of neocortical neurons to temporally modulated noisy inputs in vitro. Cerebral cortex, 18(9), pp.2086–2097.

Lam, M. et al., 2017. Large-Scale Cognitive GWAS Meta-Analysis Reveals Tissue-Specific Neural Expression and Potential Nootropic Drug Targets. Cell reports, 21(9), pp.2597–2613.

McDaniel, M., 2005. Big-brained people are smarter: A meta-analysis of the relationship between in vivo brain volume and intelligence. Intelligence, 33(4), pp.337–346.

Mohan, H. et al., 2015. Dendritic and Axonal Architecture of Individual Pyramidal Neurons across Layers of Adult Human Neocortex. Cerebral cortex, p.bhv188.

Narr, K.L. et al., 2007. Relationships between IQ and regional cortical gray matter thickness in healthy adults. Cerebral cortex, 17(9), pp.2163–2171.

Nemenman, I. et al., 2008. Neural coding of natural stimuli: information at sub-millisecond resolution. K. J. Friston, ed. PLoS computational biology, 4(3), p.e1000025.

Posthuma, D. et al., 2002. The association between brain volume and intelligence is of genetic origin. Nature neuroscience, 5(2), pp.83–84.

Salinas, E. & Sejnowski, T.J., 2001. Correlated neuronal activity and the flow of neural information. Nature reviews. Neuroscience, 2(8), pp.539–550.

Scholtens, L.H. et al., 2014. Linking macroscale graph analytical organization to microscale neuroarchitectonics in the macaque connectome. The Journal of neuroscience, 34(36), pp.12192–12205.

Sniekers, S. et al., 2017. Genome-wide association meta-analysis of 78,308 individuals identifies new loci and genes influencing human intelligence. Nature genetics, 11, p.11.

Testa-Silva, G. et al., 2014. High bandwidth synaptic communication and frequency tracking in human neocortex. I. Segev, ed. PLoS biology, 12(11), p.e1002007.

Testa-Silva, G. et al., 2010. Human synapses show a wide temporal window for spike-timing-dependent plasticity. Frontiers in synaptic neuroscience, 2, p.2.

Trampush, J.W. et al., 2017. GWAS meta-analysis reveals novel loci and genetic correlates for general cognitive function: a report from the COGENT consortium. Molecular psychiatry, 22(3), pp.336–345.

van den Heuvel, M.P. et al., 2015. Bridging Cytoarchitectonics and Connectomics in Human Cerebral Cortex. The Journal of neuroscience, 35(41), pp.13943–13948.

Verhoog, M.B. et al., 2016. Layer-specific cholinergic control of human and mouse cortical synaptic plasticity. Nature communications, 7, p.7.

Verhoog, M.B. et al., 2013. Mechanisms underlying the rules for associative plasticity at adult human neocortical synapses. The Journal of neuroscience, 33(43), pp.17197–17208.

Vernon, P., 1983. Speed of information processing and general intelligence. Intelligence, 7(1), pp.53–70.

Vetter, P., Roth, A. & Häusser, M., 2001. Propagation of action potentials in dendrites depends on dendritic morphology. Journal of neurophysiology, 85(2), pp.926–937.

